## Supplementary material for "Unbend: Correction of local beam-induced sample motion in cryo-EM images using a 3D spline model": Figure supplements

Figure 1—figure supplement 1

**Algorithm S1 | Unbend movie alignment and local motion correction**

**Input:** Movie frames $\{I_f\}_{f=1}^{N_F}$ and movie frames metadata.
**Output:** Motion-corrected micrograph.

1. **Full-frame alignment:** Perform three-iteration global alignment (Unblur-style) with increasing resolution. Identify outlier frames by deviation from an SG reference trajectory (>1.5×IQR), re-align outliers with 2×B-factor, and retain raw shifts only if they remain outliers (see Full-frame alignment)
2. **Patch trimming:** Choose patch size and patch grid $(NP_x, NP_y)$ based on output pixel size or user settings; partition the globally aligned frames into patch stacks (see Patch alignment)
3. **Patch-stack alignment:** For each patch, estimate per-frame shifts $\{(x_{i,f}, y_{i,f})\}$ by cross-correlation of each frame against the leave-one-out average of the remaining frames. (see Patch alignment)
4. **Patch smoothing and repair:** Smooth each patch trajectory using an SG filter. Flag unreliable patches using inter-frame shift variability (Eqs. 1–2; 1.5×IQR) and replace their shifts with those from the nearest-neighbor patch. (Patch alignment).
5. **Spline model and knot grid:** Construct a 3D deformation model using bicubic B-splines in $x$–$y$ and cubic B-splines along exposure/frame ($z$), with free-end boundary conditions. Set $(NK_x, NK_y) \approx \frac{2}{3}(NP_x, NP_y)$ (minimum 4), place $z$-knots every $4e^-/\mathring{A}^2$, and maintain knot parameters $K_x$and $K_y$. (see Sample distortion modeling and correction: knot grid configuration)
6. **Initialize knots:** Fit $(K_x, K_y)$ by minimizing the least-squares objective $L_1$(Eq. 11). (see Sample distortion modeling and correction: Control points and knots)
7. **Refine knots:** Refine $(K_x, K_y)$ by minimizing the cross-correlation based objective $L_2$ using L-BFGS (Eqs. 12–13). (Sample distortion modeling and correction: Parameter refinement)
8. **Warp and sum:** Evaluate pixel-wise shifts, warp frames (bilinear interpolation), pad/trim boundaries, and average corrected frames to generate the final micrograph. (see Sample distortion modeling and correction: Parameter refinement)

Figure 4—figure supplement 1

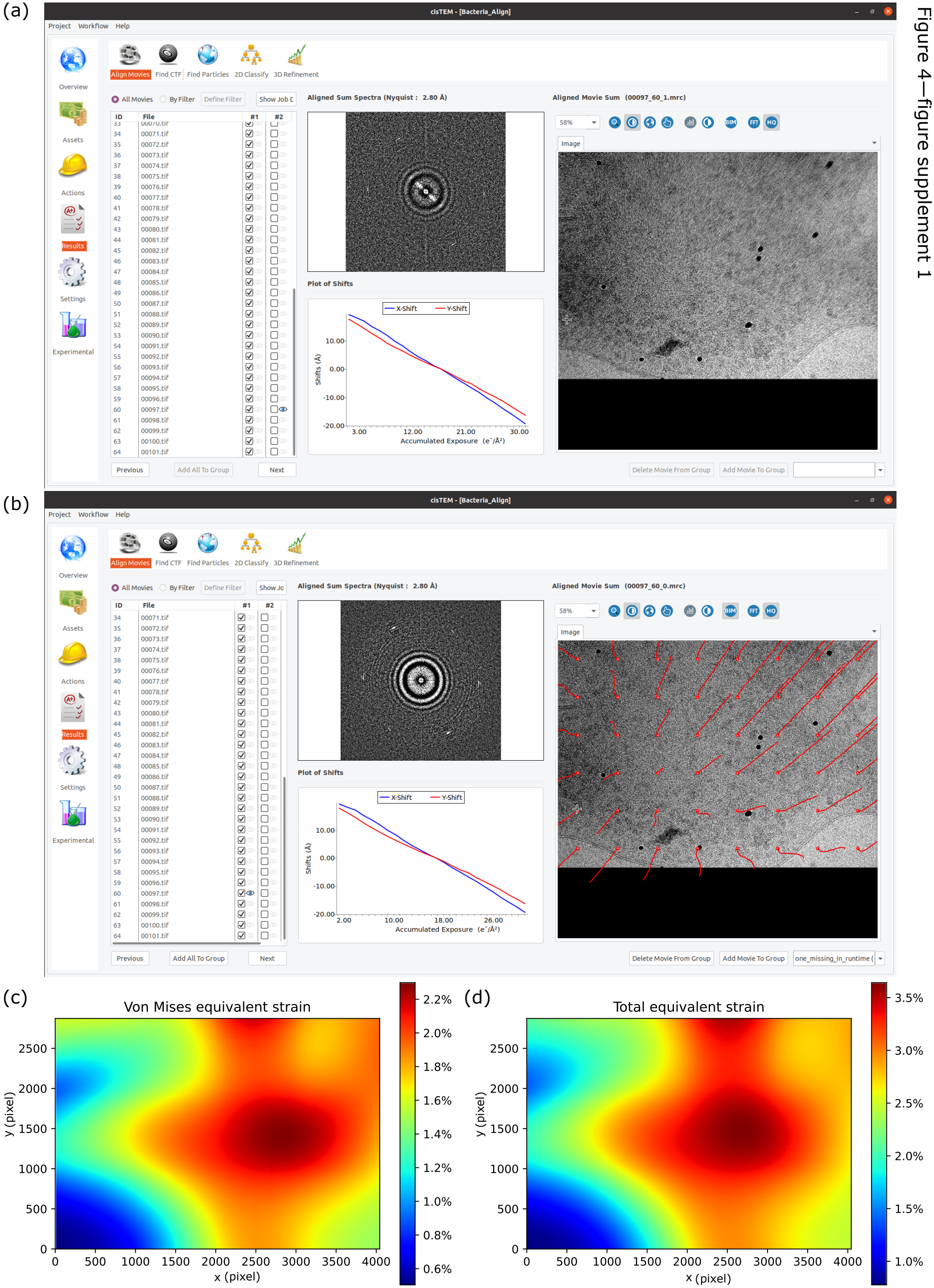

Figure 4—figure supplement 2

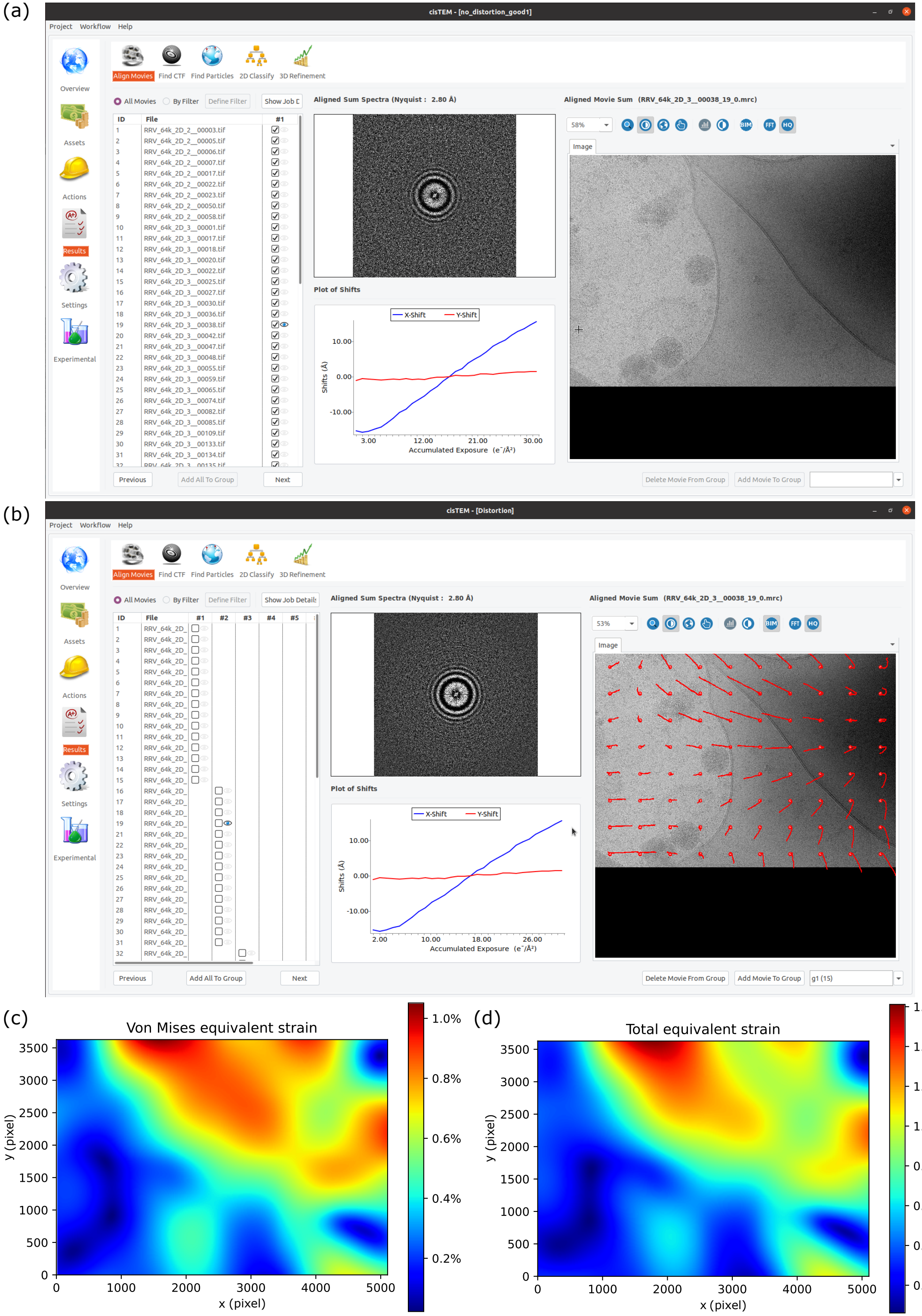

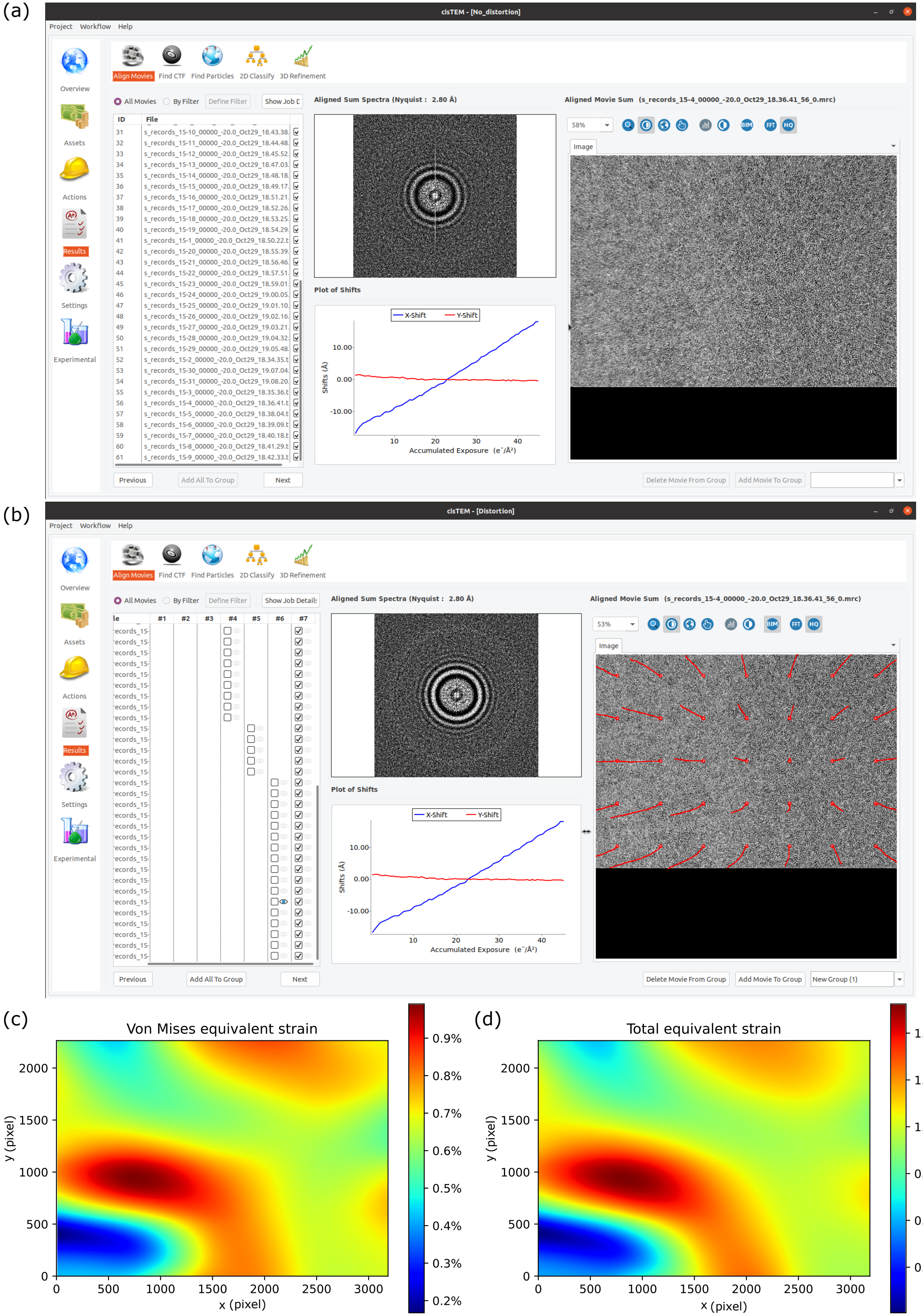


Figure 4—figure supplement 3

Figure 4—figure supplement 4

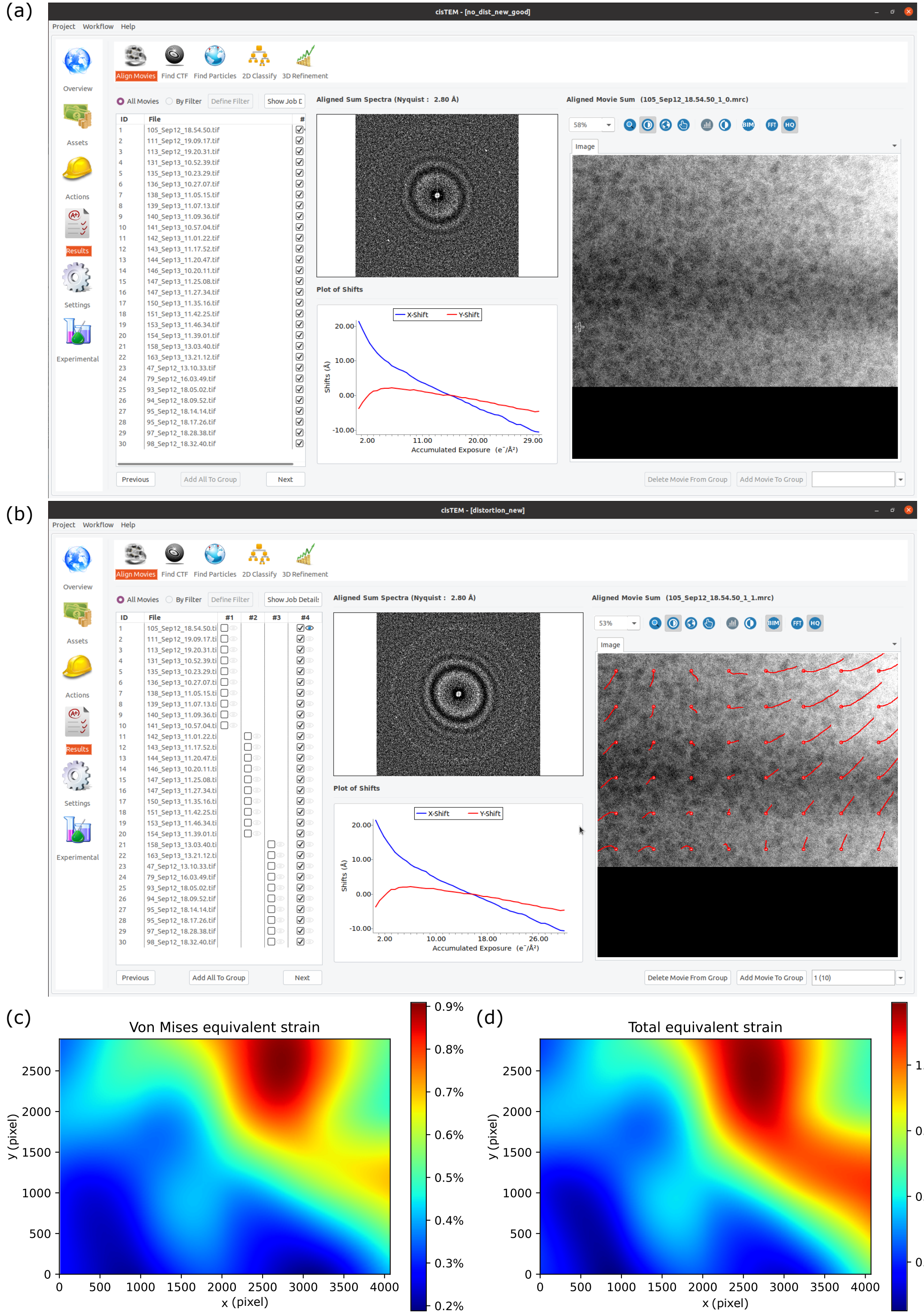

Figure 4—figure supplement 5

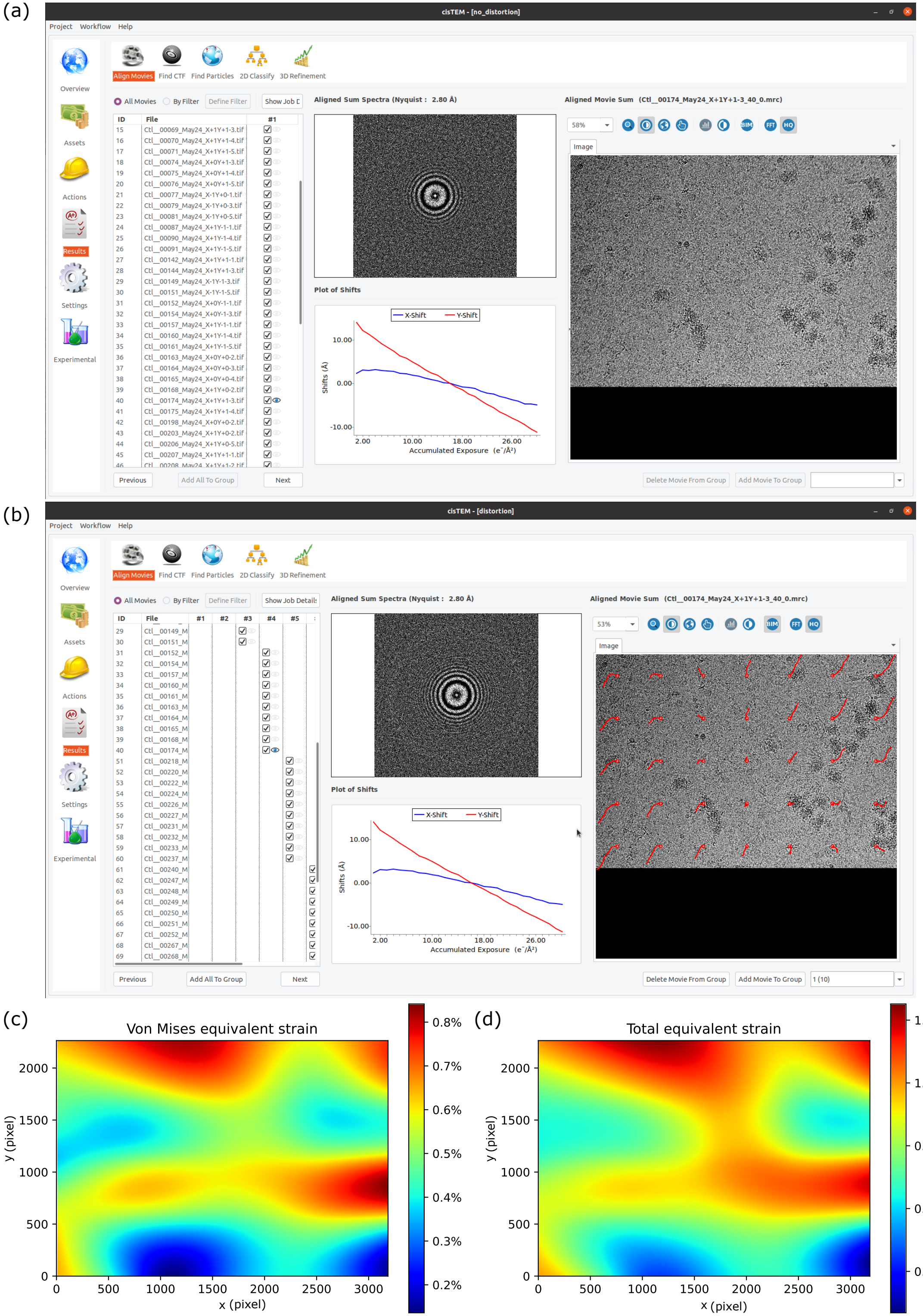

#### Figure 4—figure supplement 6 Motion and equivalent strain summary table for micrographs shown in Figure 4—figure supplements 1–5

| | Max shift (Å) | Mean shift (Å) | Standard deviation (Å) | Von Mises equivalent strain | Total equivalent strain |
|---|---|---|---|---|---|
| **M. pneumoniae** | 57.9 | 17.5 | 13.7 | < 2.3% | < 3.5 % |
| **C. aethiops: BS-C-1 (cell edge)** | 16.8 | 7.8 | 3.9 | < 1.0 % | < 1.6 % |
| **M. musculus: ER-HoxB8 lamellae** | 18.9 | 8.1 | 4.6 | < 1.0 % | < 1.5 % |
| **S. cerevisiae lamellae** | 13.2 | 5.3 | 3.6 | < 0.9 % | < 1.2 % |
| **C. aethiops: BS-C-1 (cell lysate)** | 9.1 | 3.4 | 2.3 | < 0.8 % | < 1.2 % |

For each micrograph, the mean patch shift was computed across patches; the table reports summary statistics of these per-micrograph means for each dataset.

### Figure 4—figure supplement 7 Summary statistics of per-micrograph mean patch shifts by sample type.

| Sample | Mean (Å) | Standard deviation (Å) | Min (Å) | Max (Å) | Median (Å) |
|---|---|---|---|---|---|
| **M. pneumoniae** | 4.31 | 2.17 | 0.27 | 17.46 | 4.10 |
| **BS-C-1 (cell edge)** | 2.45 | 1.70 | 0.39 | 7.78 | 2.35 |
| **S. cerevisiae (lamella)** | 1.44 | 1.04 | 0.28 | 5.30 | 1.24 |
| **ER-HoxB8 (lamella)** | 2.48 | 1.68 | 0.50 | 8.10 | 2.04 |
| **BS-C-1 (cell lysate)** | 0.72 | 0.67 | 0.29 | 3.44 | 0.49 |

#### Figure 8—figure supplement 1

#### Statistics table for detected particles from micrographs processed by different software

| Method | 2DTM SNR (mean) | 2DTM SNR (median) | Detections per micrograph (mean) | Detections per micrograph (median) | Total detections | Common detections with Unbend | Fraction of overlap (Unbend having higher SNR) (%) | Fraction of overlap (Unbend having lower SNR) (%) | Fraction of overlap (Unbend having same SNR) (%) |
|---|---|---|---|---|---|---|---|---|---|
| **M. pneumoniae** | | | | | | | | | |
| **Unbend** | 8.72 | 8.53 | 79.7 | 81.0 | 4864 | | | | |
| **MotionCor2** | 8.70 | 8.52 | 70.6 | 73.0 | 4450 | 3793 | 53.2 | 46.1 | 0.7 |
| **MotionCor3** | 8.67 | 8.47 | 67.2 | 69.5 | 4167 | 3559 | 57.1 | 42.1 | 0.8 |
| **Warp** | 8.64 | 8.47 | 60.7 | 57.5 | 3761 | 3217 | 62.2 | 37.0 | 0.8 |
| **CryoSPARC** | 8.65 | 8.45 | 50.4 | 49.0 | 3025 | 2410 | 59.1 | 40.1 | 0.8 |
| **BS-C-1 (cell edge)** | | | | | | | | | |
| **Unbend** | 10.46 | 10.26 | 70.6 | 53.5 | 4521 | | | | |
| **MotionCor2** | 10.18 | 9.98 | 67.8 | 50.5 | 4336 | 4191 | 71.6 | 28.1 | 0.4 |
| **MotionCor3** | 10.14 | 9.94 | 68.1 | 53.0 | 4293 | 4162 | 74.3 | 25.3 | 0.4 |
| **Warp** | 10.20 | 10.00 | 67.6 | 50.0 | 4324 | 4180 | 70.6 | 28.6 | 0.8 |
| **CryoSPARC** | 10.08 | 9.85 | 65.6 | 50.5 | 4197 | 3931 | 73.5 | 25.9 | 0.5 |
| **S. Cerevisiae (lamella)** | | | | | | | | | |
| **Unbend** | 10.65 | 10.51 | 407.4 | 408.5 | 12223 | | | | |
| **MotionCor2** | 10.63 | 10.48 | 406.4 | 406.0 | 12192 | 11785 | 49.9 | 49.4 | 0.7 |
| **MotionCor3** | 10.52 | 10.40 | 404.1 | 411.0 | 12122 | 11741 | 54.8 | 44.6 | 0.6 |
| **Warp** | 9.85 | 9.66 | 361.4 | 347.5 | 10842 | 10664 | 89.4 | 10.4 | 0.2 |
| **CryoSPARC** | 10.69 | 10.56 | 415.6 | 413.5 | 12469 | 11436 | 44.2 | 55.4 | 0.5 |
| **ER-HoxB8 (lamella)** | | | | | | | | | |
| **Unbend** | 11.46 | 11.35 | 10.7 | 10.0 | 407 | | | | |
| **MotionCor2** | 11.27 | 11.26 | 9.7 | 8.0 | 416 | 389 | 58.9 | 40.4 | 0.8 |
| **MotionCor3** | 11.30 | 11.23 | 9.7 | 7.5 | 406 | 388 | 63.7 | 35.8 | 0.5 |
| **Warp** | 10.82 | 10.83 | 10.1 | 9.0 | 395 | 369 | 81.8 | 17.6 | 0.5 |
| **CryoSPARC** | 10.84 | 10.74 | 9.4 | 7.0 | 405 | 345 | 79.4 | 20.3 | 0.3 |
| **BS-C-1 (cell lysate)** | | | | | | | | | |
| **Unbend** | 14.24 | 14.43 | 38.8 | 28.0 | 2910 | | | | |
| **MotionCor2** | 14.12 | 14.30 | 38.8 | 28.0 | 2909 | 2892 | 58.4 | 40.5 | 1.0 |
| **MotionCor3** | 14.11 | 14.28 | 38.8 | 28.0 | 2911 | 2890 | 58.7 | 40.6 | 0.8 |
| **Warp** | 14.02 | 14.21 | 38.8 | 28.0 | 2909 | 2887 | 62.8 | 36.7 | 0.5 |
| **CryoSPARC** | 13.71 | 13.93 | 38.1 | 28.0 | 2899 | 2875 | 72.7 | 26.9 | 0.5 |
